# The shape and volume of air, kernels and cracks, in a nutshell

**DOI:** 10.1101/2023.09.26.559651

**Authors:** Erik J. Amézquita, Michelle Y. Quigley, Patrick Brown, Elizabeth Munch, Daniel H. Chitwood

## Abstract

Walnuts are the second most produced and consumed tree nut, with over 2.6 million metric tons produced in the 2022-23 harvest cycle alone. The United States is the second largest producer, accounting for 25% of the total global supply. Nonetheless, producers face an ever-growing demand in a more uncertain climate landscape, which requires effective and efficient walnut selection and breeding of new cultivars with increased kernel content and easy-to-open shells. Past and current efforts select for these traits using hand-held calipers and eye-based evaluations. Yet there is plenty of morphology that meets the eye but goes unmeasured, such as the volume of inner air or the convexity of the kernel. Here, we study the shape of walnut fruits based on X-ray CT (Computed Tomography) 3D reconstructions. We compute 49 different morphological phenotypes for 1264 individuals comprising 149 accessions. These phenotypes are complemented by traits of breeding interest such as ease of kernel removal and kernel weight. Through allometric relationships —relative growth of one tissue to another—, we identify possible biophysical constraints at play during development. We explore multiple correlations between all morphological and commercial traits, and identify which morphological traits can explain the most variability of commercial traits. We show that using only volume and thickness-based traits, especially inner air content, we can successfully encode several of the commercial traits.

**Core Ideas:**

- X-ray Computed Tomography (CT) imaging is used to compute a broad array of morpho-logical phenotypes in walnuts.
- These morphological traits suggest biophysical constraints at play during walnut development.
- Relative inner air, shell, and packing tissue volumes are significantly correlated to the rest of shape phenotypes.
- These volumes produce the best prediction models for traits of commercial interest such as shell strength.
- Inexpensive phenotyping platforms that focus solely on volume measurement would enable better walnut breeding.

## 1 Introduction

There is more than meets the eye in the shape of walnuts (*Juglans regia*). Civilizations originary from modern day Iran have used and traded walnut fruit and tree products since the 7000 BC (Vahdati, 2014). From there and then, walnuts traveled far and wide as they were actively traded through the Silk Road, conquering the Eurasian continent (Pollegioni et al., 2014). The trade of walnuts remains an important part of the global economy. In 2021, the California and the US produced more than 725,000 tons of walnuts valued in more than $1.0B, following a historically increasing trend of both bearing acreage and bearing trees per acre (NASS, 2022). World demand for walnut keeps increasing and it is estimated that the world will consume a record 2.5M tons of walnuts for 2023, and the US is forecasted to satisfy 25% of the global demand (FAS, 2022). The trade is not limited to the food industry, as there is also growing research on additional uses for walnut shell material. This material can be key for more durable batteries (Wahid et al., 2017), lower-cost concrete (Hilal et al., 2020), and stronger epoxy composites (Lala et al., 2018), to name a few examples. As climate change alters weather patterns, and the demand for walnut and its byproducts increases, we must breed walnuts with more suitable traits such as high kernel-to-total weight ratio, adequate shell strength, and easiness to extract both kernel halves intact. Quantitative analyses and comprehensive pheontyping can accelerate current breeding programs by quickly identifying varieties and individuals with desirable characteristics (Fiorani and Schurr, 2013; Rahaman et al., 2015). The rapid selection of potentially desirable progenitors for breeding programs is especially crucial for walnuts, as seedlings are hard to propagate. Even for the fastest-growing accessions, it takes at least 2 years for trees to bear fruit for the first time, and at least 5 more to yield fruit at a commercial scale (Lopez, 2004; Popa et al., 2023).

Most of the current walnut phenotyping follows the measuring guidelines set by the International Plant Genetic Resources Institute (IPGRI, 1994). The morphological phenotyping of the fruit is mainly done using calipers to measure length, width, and height, combined with visual assessments to describe more complicated traits such as texture and sphericity. These simple measurements have proved to be insightful to evaluate and identify promising genotypes. Moderate correlations have been reported between these traditional morphological traits of the walnut tree and fruit with commercial and horticultural traits of interest such as pollen release strategy, yield, shell thickness, kernel weight, and pathogen resistance (Akca and Şen, 1995; Kelc et al., 2007; Khadivi-Khub et al., 2015; Rezaei et al., 2018; Shah et al., 2021; Solar et al., 2003).

However, this caliper- and eye-based approach is time consuming, prone to human error and subjectivity, and fails to capture richer shape nuance observed in the shells and kernels. As next-generation sequencing (NGS) technology advances, we observe an explosion in genomics data collection that must be matched by equally powerful and encompassing phenomics (Araus and Cairns, 2014; Bucksch et al., 2017). We have to look deeper than just nut lengths and widths. To that end, X-ray computed tomography (CT) scanning has proved to be a powerful tool to accurately capture intricate, internal features of a vast array of plant data in a nondestrutive manner. High-resolution, X-ray CT 3D reconstructions have been successfully used to capture and quantify the complex branching architecture of inflorescence in grapevines (Li et al., 2019) and sorghum panicles (Li et al., 2020), identify key morphological traits in barley seeds to distinguish their accession of origin (Amézquita et al., 2021), and determine nuances in soil porosity for diverse wheat root-soil interactions (Zhou et al., 2020)

To the best of our knowledge, Bernard et al. (2020) is the first study that exploits X-ray CT imaging to automatically, accurately, and systematically quantify multiple walnut shape phenotypes. Their results showcase the morphological variability found across the germplasm diversity panel maintained by INRIA, France. There, Bernard et al. measure the absolute and relative volumes of the whole nut and its shell, kernel, and internal air. In particular, they observe that larger fruits are correlated with rougher shell shape and smaller kernel filling ratio, which allows them to select for better genotypes. X-ray CT imaging has also been recently used to document morphological changes of walnut flower bud development (Gao, 2022), estimate kernel weight in an nondestructive way (Gao et al., 2022), and to explore the puzzling diversity and structure of the cell tesselations that conform the hard shell tissue for multiple nuts (Huss et al., 2020).

Here, we study the shape of walnut fruits based on the X-ray CT 3D reconstruction of 1264 individuals comprising 149 accessions maintained by the Walnut Improvement Program at the University of California Davis. We exploit the nondestructiveness of X-rays to isolate individual walnuts and segment out shell, kernel, and packing tissues, as well as the air contained inside every walnut. We first compute 49 different shape- and size-related traits for each walnut such as nut and kernel dimensions, surface areas, volumes, filling ratios, sphericity and convexity indices. We included the computation of the 14 traits used by Bernard et al. (2020). All the image processing tasks were done with an in-house, python-based, open-source script. This morphological information was combined with values of breeding interest that were collected separately throughout different years. These traits include ease of removing both kernel halves intact, shell strength, and kernel-to-nut weight ratio. Second, we look for allometric relationships of interest across the whole population —the growth rate of a tissue relative to another— which reveal possible biophysical constraints at play during walnut development. These allometric relationships pose theoretical limits on minimum and maximum possible walnut sizes. Third, we examine Spearman correlation coefficients between all the computed morphological phenotypes. We find that the relative content of inner air, an often overlooked trait, is significantly correlated to the rest of tissue sizes and shapes. This suggests that the inner air content plays an important role during tissue development. The air volume is also significantly correlated with several traits of breeding interest. Fourth, we compute several stepwise linear regression models to determine which shape and size phenotypes contribute the most to relative kernel weight, ease of kernel removal, and shell strength. Our results suggest that these traits of breeding interest can be best predicted with only a handful of independent phenotypes, including inner air content. Finally, we examined the distribution of these phenotypes within each accession and through the general population. A principal component analysis using only the 6 most relevant morphological phenotypes reveals that accessions follow a gradient according to their shell strength scores and kernel weight ratio. This suggests that key traits can be successfully predicted if the volume of different tissues is known, which opens the possibility of engineering affordable phenotyping plantforms that only focus on volumetric analyses rather than expensive CT setups. This morphological modeling will allow us to set a new exciting path to explore further the phenotype-genotype relationship in walnuts.

## 2 Materials and methods

### 2.1 Plant material and scanning

All plant materials represent walnut breeding lines, germplasm, and cultivars maintained by the Walnut Improvement Program at the University of California, Davis. A total of 150 walnuts accessions were harvested into mesh bags at hull split, oven-dried overnight at 95°F, and then air-dried for several weeks before moving into cold storage at 35°F. 5 to 16 individuals were selected for each accession, for a total of 1301 individual walnuts to be scanned at Michigan State University (Table S1). The walnuts were scanned in 171 batches. The scans were produced using the the North Star X3000 system and the included efX-DR software. The X-ray source was set at 75 kV and 100 µA, with 720 projections per scan, at 3 frames per second and with 3 frames averaged per projection. The data was obtained in continuous mode. The 3D X-ray CT reconstruction was computed with the efX-CT software, obtaining voxel-based images with voxel size of 75.9 µm.

All the individual walnuts were manually separated with ImageJ (Figure 1A). Densities were rescaled so that all scans share similar air, kernel, and shell density values. Once densities were comparable across samples, the external air and other debris was removed through thresholding and mathematical morphology operations (Figure 1B). Rough estimates for the location of shell, air, kernel, and packing tissues were obtained based on density, connectedness, and object thickness information. These estimates were used then to fully segment the tissues using a watershed segmentation algorithm (Falcao et al., 2004) (Figure 1C-H). We took particular care of tissue labeled as shell, where we distinguished voxels close to the walnut surface, to voxels protruding into the internal cavity (Figure 1D). Some of the scanned walnuts contained incomplete or no kernel at all. These were discarded from further morphological analysis, leaving us with a total of 1264 individual walnuts representing 149 accessions (Table S1). All the image processing above was done automatically with in-house, scipy-based, python scripts.

**FIGURE 1.**
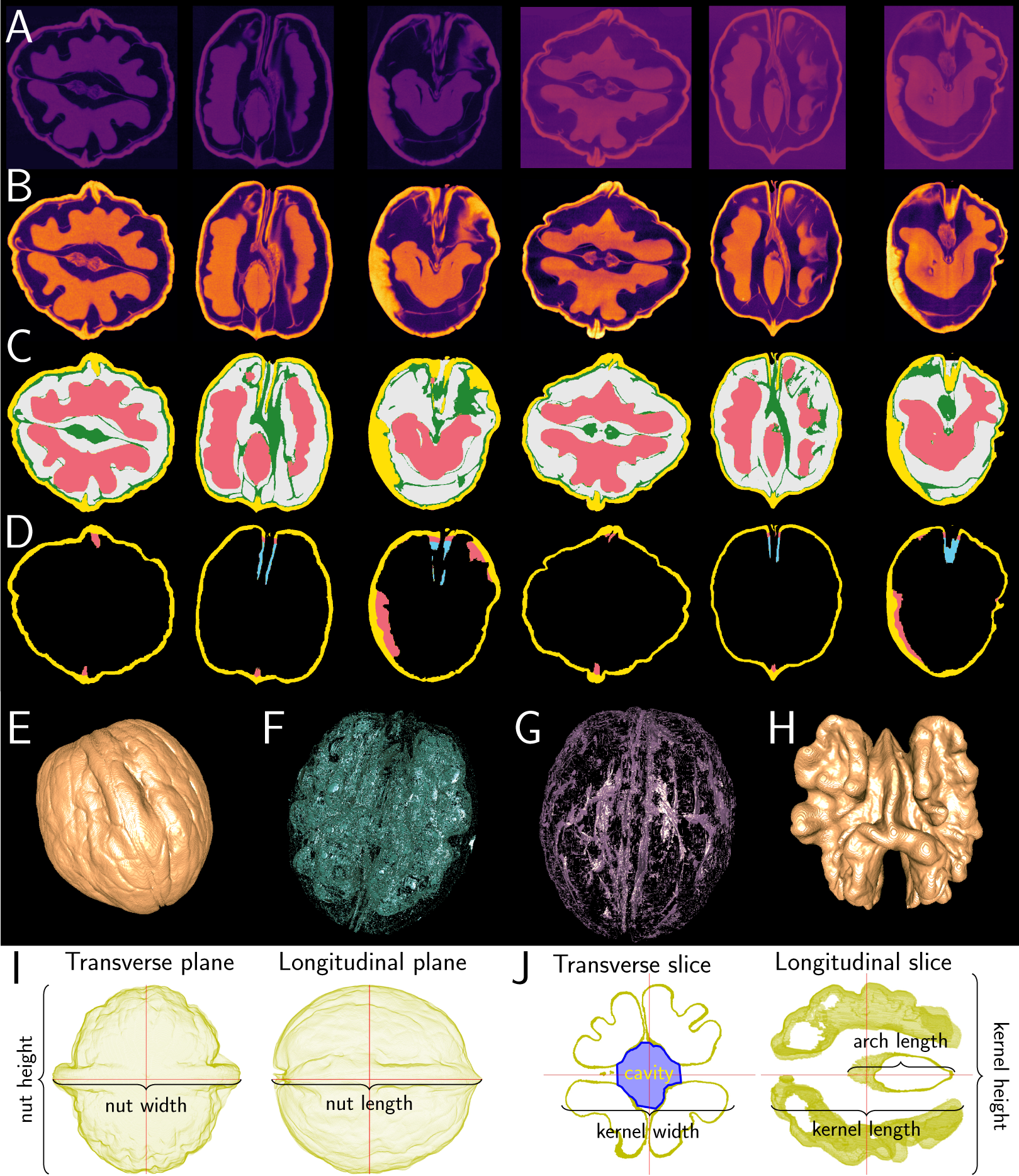
Walnut scanning, image processing, and phenotyping. (A) Raw scans of individual walnuts seen from different planes. (B) Densities were standardized across all samples and the external air removed. (C) Shell, air, kernel, and packing tissue were automatically labeled with a combination of basic image morphology operations and watershed segmentation. (D) The tissue labeled as shell was further broken down into external and protruding tissue. (E) 3D renders of shell, (F) air, (G) packing tissue, and (H) kernel. (I) All the walnuts were centered on their center of mass and aligned. (J) The same centering and alignment was applied to the kernels. All the figures above are for illustration purposes only and are not scaled.

To make some measurements comparable, all the walnuts were centered on their centers of mass and rotated such that the lateral plane goes through the walnut seal, and the shell tip is the rightmost point of the longitudinal plane (Figure 1I.) The same center and rotation was immediately applied to the kernel. By taking a series of 2D transverse slices across the proximal-distal axis, we approximated the main cavity surrounded by the kernel, located a the proximal side of the walnut, between the two main hemispheres of the kernel (Figure 1J).

### 2.2 Walnut morphological trait measurements

For each individual we computed the same 14 morphological traits as in Bernard et al. (2020): nut length, height, width, total surface area, total volume, rugosity, sphericity, shape VA3D, equancy, shell volume, shell thickness, kernel volume, kernel volume filling ratio, and the inner air volume. We computed an additional collection of 35 morphological traits for a total of 49 measurements per sample. This includes the volume of packing tissue, as well as the percentage of air, kernel, shell, and packing tissue volume with respect to the total nut volume —the sum of air, kernel, shell, and packing tissue volumes. We also considered the airless volume ratio —where now the total nut volume is just the sum of kernel, shell, and packing tissue volumes. Sphericity was calculated with the Krumbein and Sneed indices for every walnut —indices between 0 and 1, where 1 indicates a perfect sphere (Blott and Pye, 2008). The surface area and volume of the nut’s convex hull was compared to the actual nut surface area and volume respectively as a proxy for different kinds of lobeyness —the ratios are between 0 and 1, where 1 indicates a perfectly convex shape (Figure 2). These computations were repeated for the kernel to assess its sphericity and lobeyness as well. For the main cavity at the proximal side of the kernel, we measured its depth, surface area, and volume. We computed the percentage of cavity area and volume comprised by the total kernel surface area and kernel’s convex hull volume respectively as a proxy for relative size of the cavity. We also measured the length of the arch-like structure of the kernel that bridges the two main hemispheres. As indicated before, for the shell and tissue with shell-like density, we computed both the volume and the percentage of it protruding into the main walnut cavity. Finally, we computed the average density value for shell, kernel, and packing tissues. Since the X-ray CT scans only measure relative density, we considered the three possible ratios of these averages as a proxy for absolute density. (Table 1).

**FIGURE 2.**
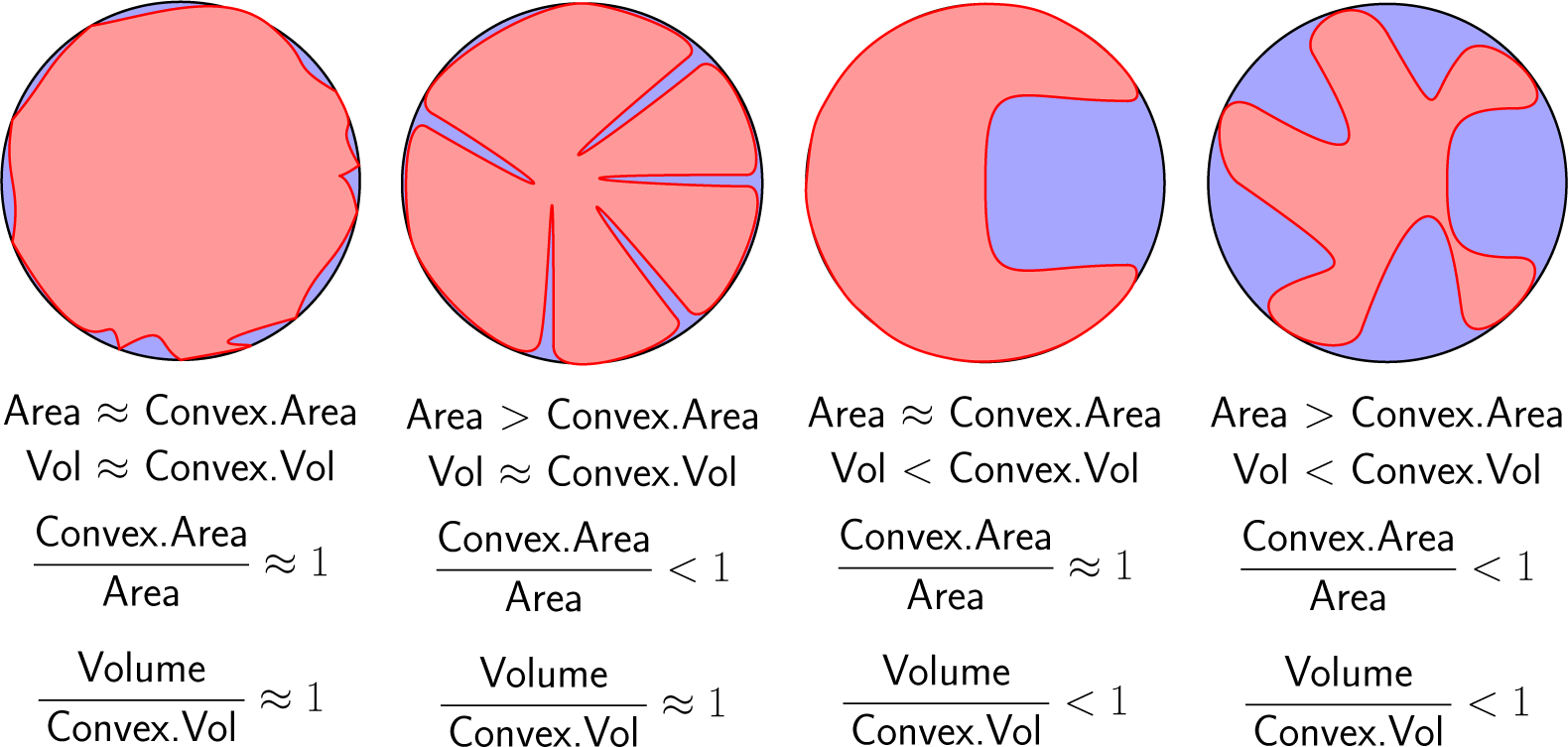
Comparing the surface area and volume of an object to the ones from its convex hull can reveal different kinds of lobeyness. For illustrative purposes, a 2D example is presented below, with a red object and its blue convex hull. As this is a 2D example instead of a 3D one, area refers to perimeter, while volume refers to area instead. Both convex area and volume ratios are bounded between 0 and 1, where 1 indicates perfectly convex shape. A low convex area ratio with a higher convex volume ratio suggests deep, narrow troughs across the object’s surface. The opposite case suggests wider, square-like cavities.

**TABLE 1.**
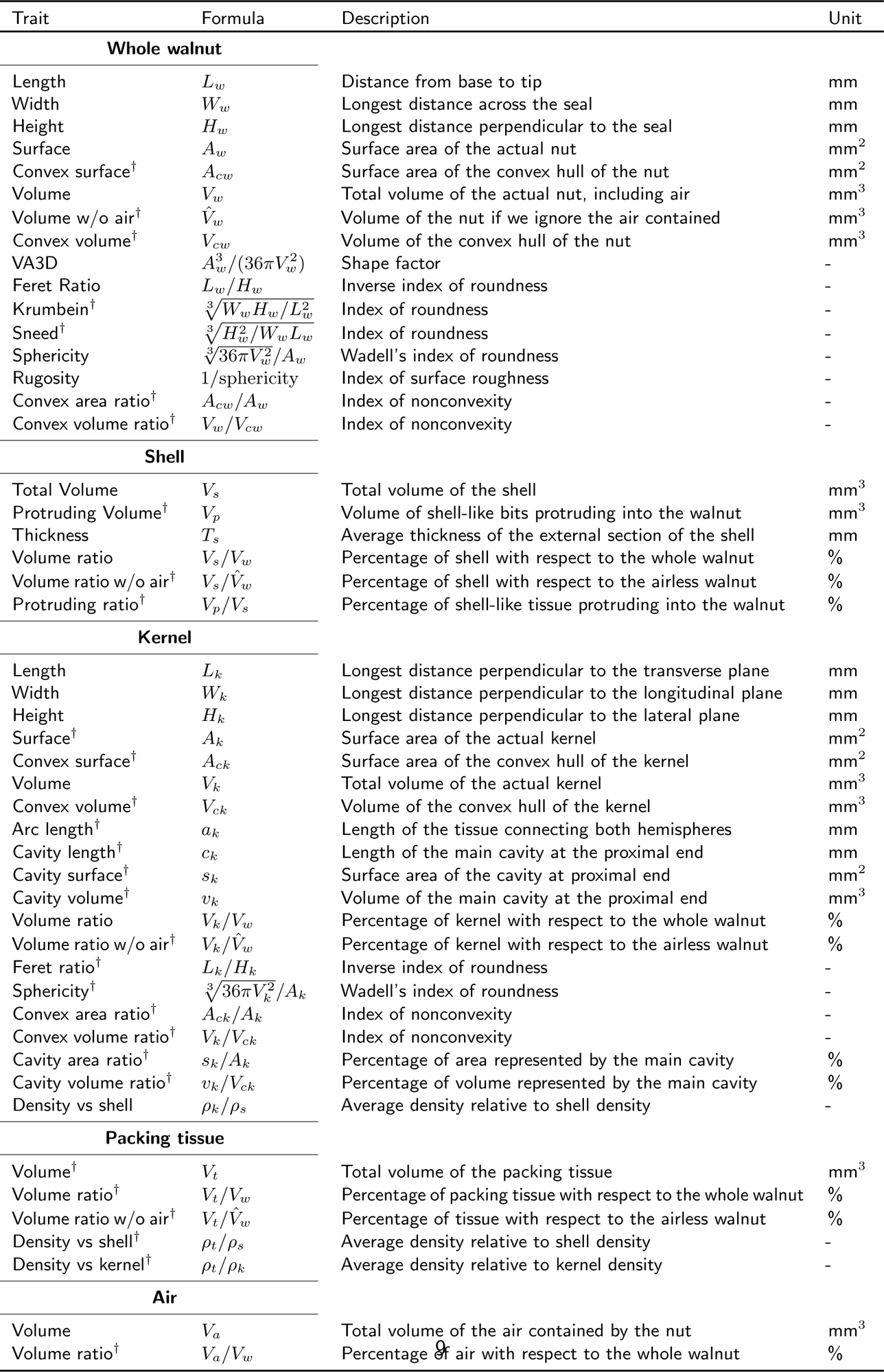
Morphological traits measured for each walnut based on its X-ray CT scan. A dash denotes not applicable. A dagger (†) denotes a trait that has not been measured before to the best of our knowledge.

### 2.3 Measurement of additional traits of breeding interest

Separately from the walnuts scanned at Michigan State University, accessions were evaluated for 14 traits of breeding interest (Table 2). Ten walnuts per accession were collected across multiple years and locations managed by the Walnut Improvement Program at the University of California, Davis. These walnuts were later cracked open from each sample using a hammer at Davis, California. The assessment and scoring was done following the IPGRI (1994) guidelines. All the trait scores were averaged across all the samples of the same accession to obtain scores representative of said accession.

**TABLE 2.**
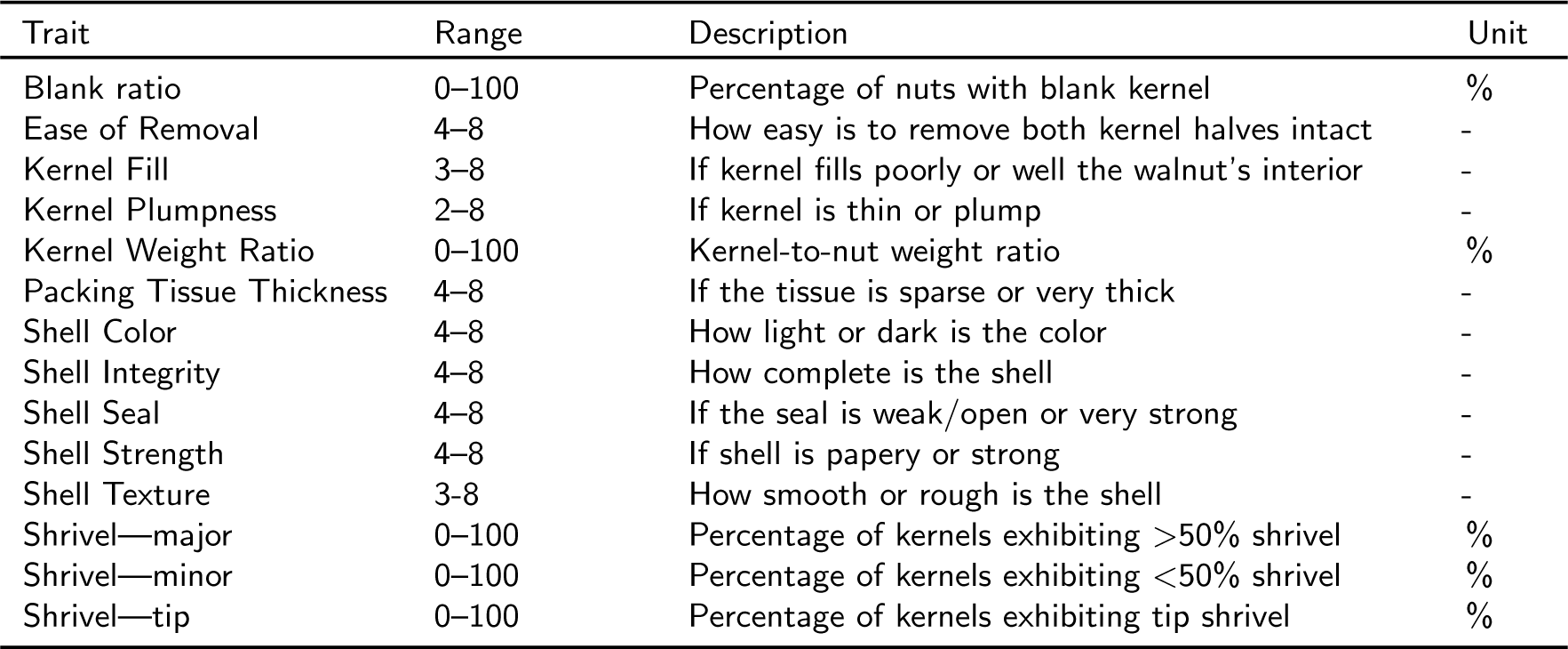
Traits of breeding interest. Measured and evaluated per IPGRI (1994) guidelines. Dash indicates not applicable.

### 2.4 Allometric relationships

We measured the variability of each morphological phenotype by computing their quartile coefficient of variation (QCD) across the 1264 scans. We preferred the QCD as it only depends on the 25th and 75th quartiles, making it robust against outliers compared to the coefficient of variation (CV) (Bonett, 2006). We studied allometric relationships between all size-specific morphological traits —the relative growth of one feature with respect to another one. Since different plant tissues grow relative to each other following a power law rather than a simple linear relationship (Niklas, 2004; West et al., 1999), we plotted size-related data in log-log plots and calculated the best fit line (Figure 3). The slope of this line indicates whether a tissue grows at a faster or slower pace than another, while the intercept allows to compute theoretical limits on tissue size.

**FIGURE 3.**
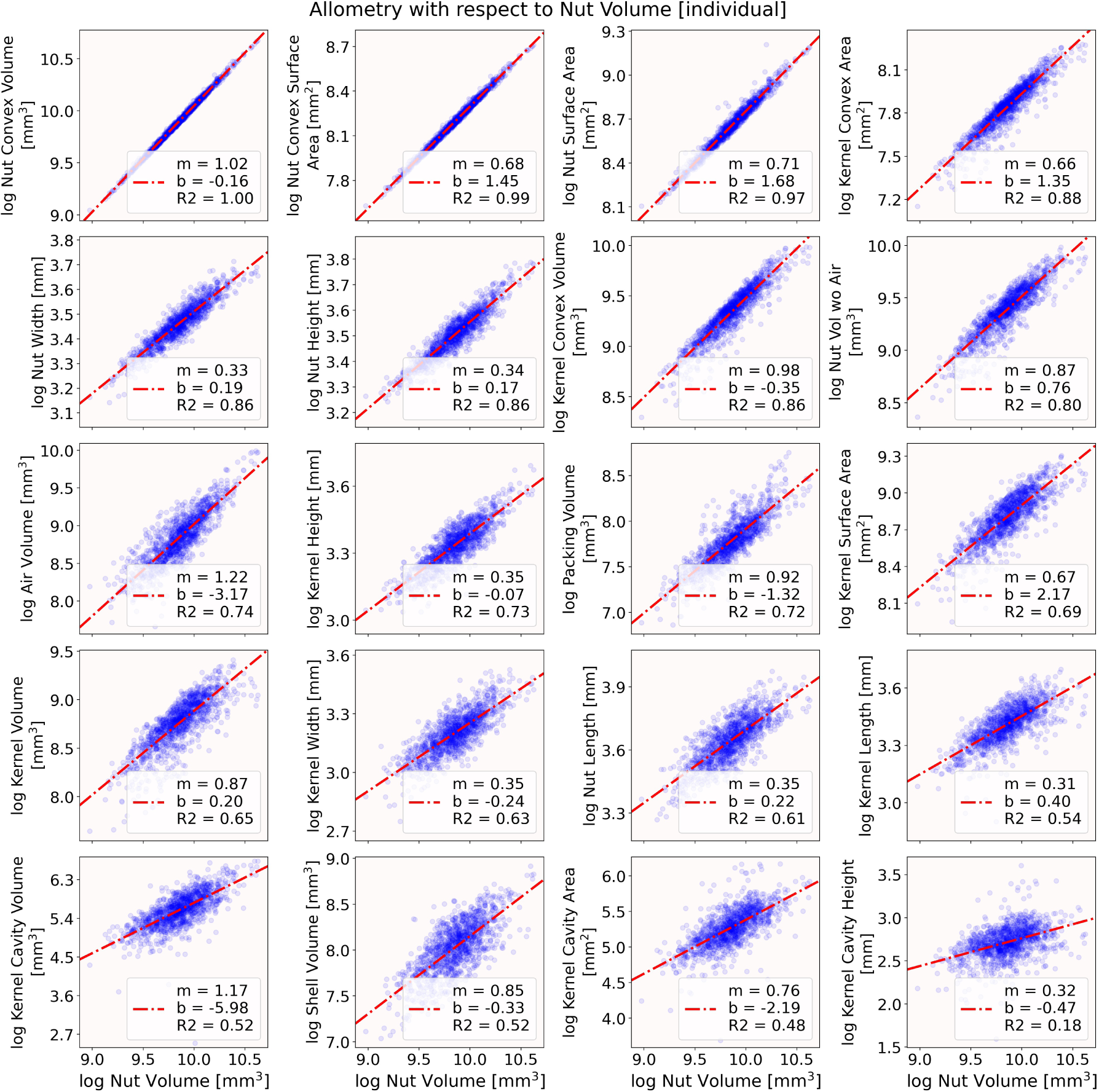
Various allometry plots between different logarithmic values of tissue volumes, areas, and lengths compared to the total walnut volume. An ordinary least squares linear model was computed for each case. The slope, intercept, and coefficient of determination for each linear model is indicated by *m*, *b*, and *R*^2^ respectively.

### 2.5 Accession summaries and correlations

We averaged each morphological phenotype for walnuts within the same accession so we could compare them to the accession-based traits of breeding interest. Since size-related phenotypes follow nonlinear relationships, we favored the computation of Spearman rather Pearson correlation coefficients between all the different morphological and breeding phenotypes (Figures 4, S1, S2). After applying a Bonferroni correction, all p-values smaller than 0.01 were deemed highly significant.

**FIGURE 4.**
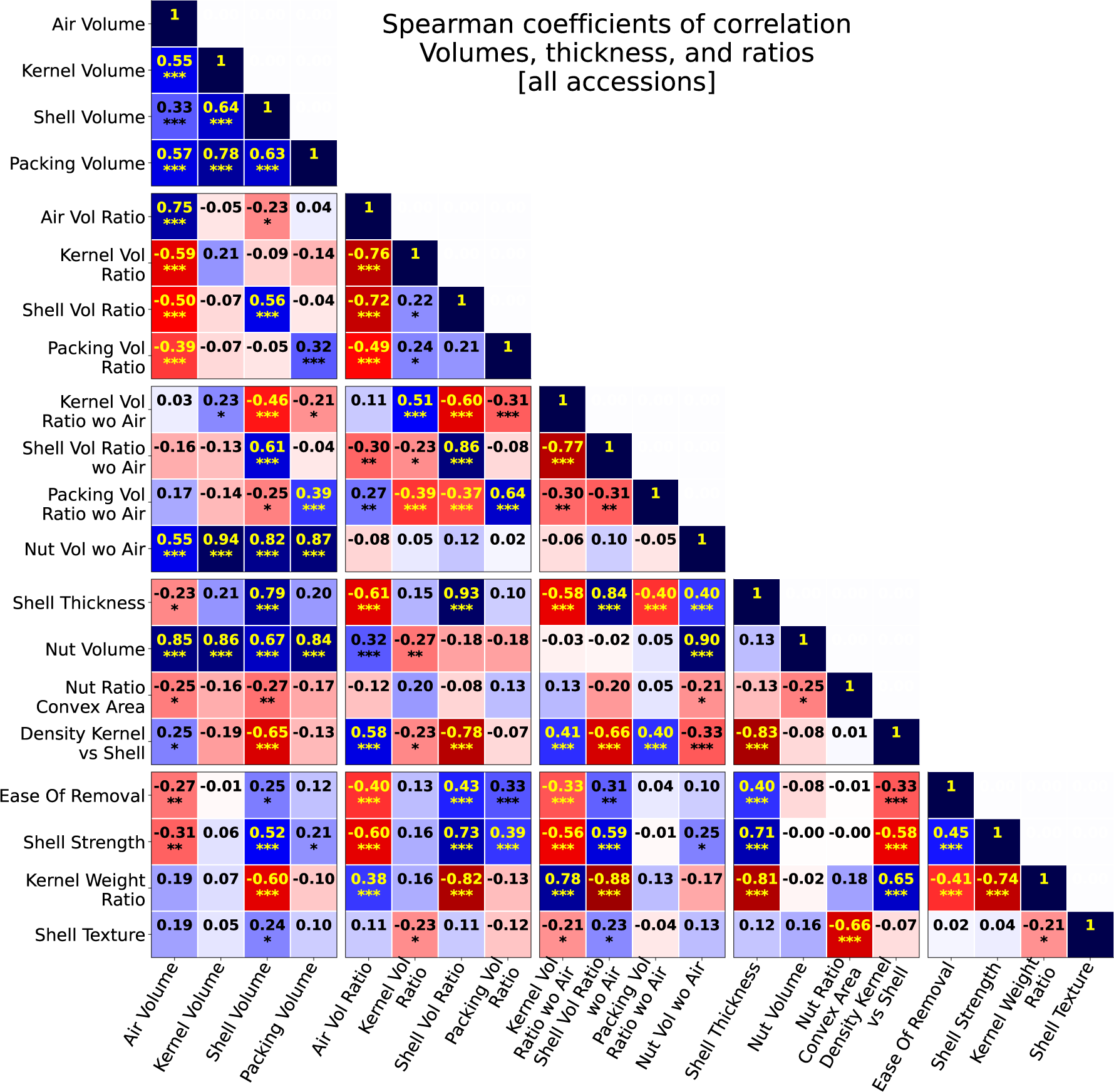
Spearman correlation coefficients between shape, size, and traits of breeding interest. The correlation was computed using phenotype values that were averaged across all individuals of the same accession. Three stars (***) denote an associated p-value of 10*^−^*^4^ or smaller. Two stars denote a p-value between 10*^−^*^4^ and 10*^−^*^3^. One star denotes it between 10*^−^*^3^ and 10*^−^*^2^.

### 2.6 Prediction of traits of breeding interest

A stepwise linear regression was performed to determine the most relevant morphological phenotypes to predict ease of removal. We only considered phenotypes significantly correlated to ease of removal as potential explanatory variables. Data was standardized to zero mean and unit variance. A train-test split was followed to avoid overfitting results. That is, an ordinary least squares (OLS) linear model was fitted using 70% of the data while predictions of ease of removal were made for the remaining 30%. The *R*^2^ coefficient of determination was computed between the predicted and actual ease of removal values as a measure of prediction accuracy. The 70-30 training-test split was randomized and repeated 100 times and the average *R*^2^ coefficient was taken as the overall accuracy score. To determine the contribution of individual phenotypes to the prediction of ease of removal, we computed numerous linear models where we varied the number and the specific phenotypes used as explanatory variables. First we computed models using a single explanatory phenotype and recorded the model with the highest accuracy. Second, we considered models using all the possible pairs of phenotypes as explanatory variables and recorded the pair that provided the model with the highest accuracy. We subsequently repeated this exploration of models using all possible combinations of three to six phenotypes as explanatory variables (Figure 5A). The above procedure was repeated to identify the morphological features that are the most relevant to predict shell strength and kernel weight ratio (Figure 5B-C). For all the models described above, we noticed that prediction results improved considerably if the Earliest Himalayan accession (UCACCSD 85-023-2) was discarded (Figure S3). All the *R*^2^ scores reported are thus considering only 148 of the 149 accessions scanned.

**FIGURE 5.**
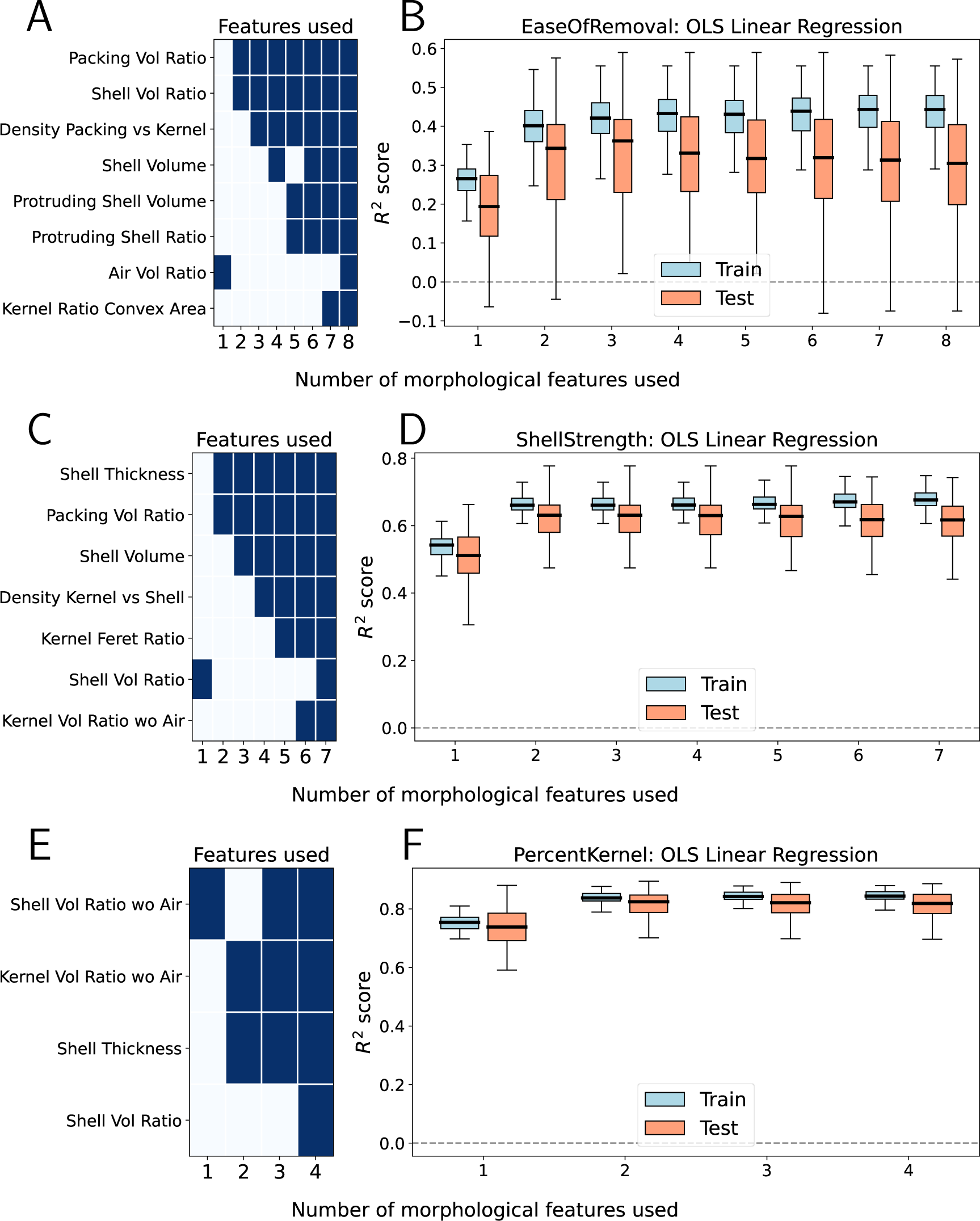
Morphological traits that can predict the best certain traits of breeding interest using an OLS linear regression. (A) If the model is limited to a single explanatory variable, then air volume ratio produces the best possible prediction of ease of kernel removal. (B) The *R*^2^ coefficient between predicted versus the true ease of removal value is 0.2. This can improve up to 0.32 if the OLS model uses two explanatory variables instead, these two being shell and packing tissue volume ratio. A moderate improvement is made by considering three explanatory variable, adding relative kernel density. No improvement is made by adding more explanatory variables. (C) - (F) Analogous analyses were made with models predicting shell strength, relative kernel weight.

Finally, we performed a principal component analysis (PCA) of the most predictive morphological phenotypes. We then plotted the two principal components and observed how these also reflect the traits of breeding interest. The PCA was repeated using all 49 morphological phenotypes as well.

## 3 Results

### 3.1 Distribution and variability of phenotypes

Values of nut length, width, height, sphericity, convexity, and density were overall stable with low QCD values (0.05 or less). This suggests that walnuts by and large have similar overall shell shape, rugosity, and shell lobeyness. A high nut convex volume ratio (0.95±0.01) combined with a low convex surface area ratio (0.63±0.01) indicates that most of the shell’ surfaces are covered by numerous, narrow, deep grooves. The relative average densities are also stable (0.03), where the reported average density across the whole kernel and packing tissue is 86% and 61% of the shell density respectively. This is contrasted by higher variability of tissue volume (0.15), which agrees with reports of highly variable walnut weight values (Cosmulescu and Stefanescu, 2018; Rashnodi et al., 2019). At the same time, walnuts reveal a especially large variability of shell tissue. In particular, the amount of shell tissue that protrudes into the walnut ranges from 12 to 1077 mm^3^. This corresponds to a QCD of almost 0.4, where the 75th percentile (237 mm^3^) is more than double of the 25th one (106 mm^3^). There is similar variability when it comes to all the measurements related to the main proximal-side cavity (Table 3).

**TABLE 3.**
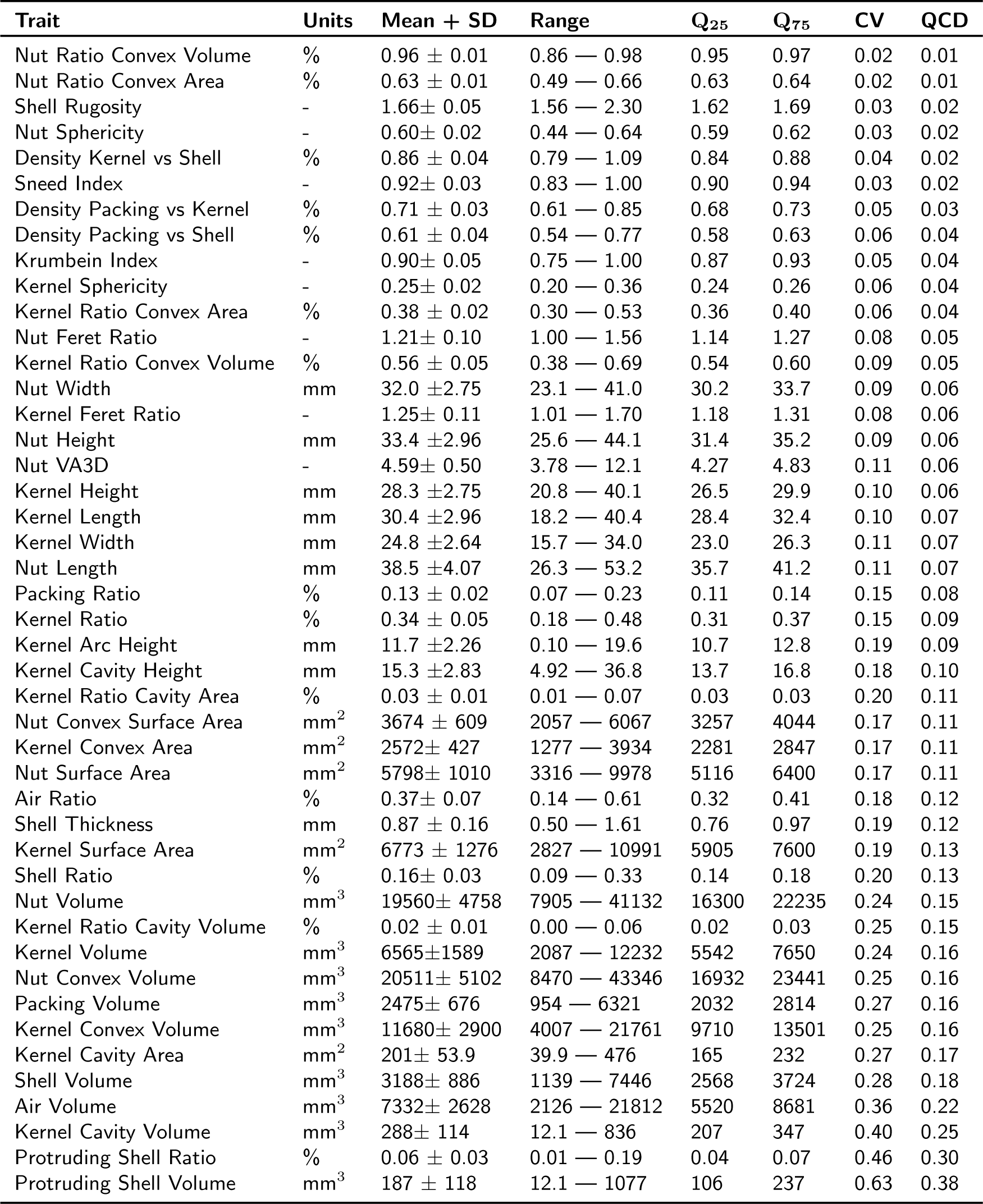
Morphological trait values. Standard deviation, 25th quartile, 75th quartile, quartile coefficient of dispersion, and coefficient of variance are indicated by SD, **Q_25_**, **Q_75_**, QCD, and CV respectively. Traits sorted by QCD. A dash denotes not applicable.

### 3.2 Allometry and possible biophysical constraints

We observe that most of the size-specific traits follow power laws with respect to the total nut volume *V_w_*, as our allometric log-log plots exhibit large *R*^2^ coefficients of determination, most of them above 0.5 (Figure 3). The size-related measurement that exhibits the most superlinear growth rate is the total air volume contained inside the nut *V_a_*. Our data suggests that these two volumes follow the power law 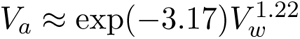. That is, as the nut volume increases, biophysical constraints require an air volume increase by a larger factor. However, the air volume must always be lower than the total nut volume. Evaluating the extreme case of a hypothetical walnut consisting entirely of air, we find that 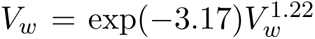 when *V_w_* ≈ 2.3 ×10^6^ mm^3^. This is the same volume of a 16cm diameter sphere. The volume of the convex hull of the nut *V_cw_*also follows a superlinear growth rate with respect to *V_w_*, with a power law 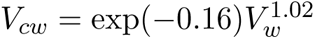. Do notice that if *V_w_ <* 2063mm^3^, then *V_cw_ < V_w_*, which is impossible.

In other words, the allometric power law only holds for walnuts comparable to a 1.6cm diameter sphere. This agrees with the fact that the walnut seed is very lobed during early developmental stages (Sartorius and Stösser, 1997). Of important note is the fact that kernel volume grows at a slightly sublinear rate with respect to total nut volume, with a power law 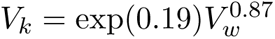. For instance, if the total nut volume is duplicated, then the kernel volume will only increase by a factor of 2^0.87^ ≈ 1.8, which already indicates that larger walnuts tend to have smaller kernel percentages, while they also tend to contain higher air percentages.

### 3.3 Correlation between several morphological phenotypes

The last observation is also supported by a significant negative Spearman correlation between the kernel and air volume ratios (-0.59), and nut volume (-0.27). (Figures 4). If we ignore the air when taking into account volume ratios, we also observe very negative correlations with shell volume, shell volume ratios, and shell thickness (between -0.46 and -0.88). For shell thickness, we observe unsurprisingly high correlations with shell volume percentage (0.93), shell total volume (0.79), and other shell-related measurements. There are negative correlations with kernel density relative to shell density (-0.83) and with air percentage (-0.61) (Figure 4). Whenever air was discarded from the volume ratio considerations, we observed that the ratios of shell, kernel, and packing tissue were significantly and negatively correlated between each other. This suggests that increasing one ratio diminishes the other two, and that these three ratios can be manipulated independently. On the other hand, whenever air was included into the volume ratio considerations, we observed no significant correlations among percentages of shell, kernel and packing tissues. However, these three were all negatively correlated to the air ratio. This suggests that the air ratio determines the volume ratio for the rest of tissues. Another interesting observation is that the volume ratio of shell and packing tissue when the air is excluded is negatively (-0.30) and positively (0.27) correlated respectively to the volume ratio of air. This might indicate that increasing the content of air also increases the content of packing tissue at the expense of shell (Figure 4). There is also significant correlation between convexity and sphericity indices (Figure S1). Finally, we note that air ratio is positively correlated (0.54) with kernel convex area ratio while negatively correlated (-0.81) with kernel convex volume ratio, which suggests that walnuts with large air content tend to have large and wide troughs (Figure S2A). However, there was no strong correlation between air ratio and the main cavity at the proximal side of the kernel (Figure S2A).

### 3.4 Correlation between morphology and traits of breeding interest

Ease of kernel removal is negatively correlated to air (-0.40) and kernel (-0.33) volume ratios, and to kernel weight (-0.41) and density (-0.33) ratios. Ease of removal is positively correlated to shell (0.43) and packing tissue (0.33) volume ratios, and to shell thickness (0.40). The shell strength unsurprisingly is positively correlated with shell volume ratio (0.73) and shell thickness (0.71). It is also negatively correlated to the volume (-0.56), weight (-0.74), and density (-0.58) ratios of the kernel, and to the air volume ratio (-0.60). Finally, we observe that weight ratio of the kernel is negatively correlated with shell volume ratio (-0.82) and thickness (-0.81). We highlight that there is no correlation (0.07) between kernel weight and kernel volume ratios whenever the latter takes air volume into consideration. The correlation improves significantly (0.78) when the volume ratio does not include air, but this is not a one-to-one relationship between kernel volume and weight. This could be explained by observing also a significant correlation (0.65) between relative kernel weight and density, suggesting that as kernel increases in size, so does in density (Figure 4). In other words, nuts with a higher relative content of air tend to have weaker shells, more relative kernel weight, and the kernel is easier to extract.

### 3.5 Modeling and explaining certain traits of breeding interest

Whenever we limited an OLS linear model to use a single explanatory variable to predict ease of kernel removal scores, the best possible prediction on average was achieved by using air volume ratio. The model in this case reported an average *R*^2^ coefficient between predicted scores versus true scores was 0.26 when predicting the same data used to train the model, and 0.18 when using test data instead. We observe that through the 100 repetitions, the *R*^2^ coefficient is highly variable, which indicates that the model is quite sensitive to the initial train-test split. This large variability remains present regardless of the number of explanatory variables used by the model, which reflects that the ease of removal of scores are not normally distributed for the accessions at hand. If the OLS model is allowed to use two explanatory variables, the average *R*^2^ coefficient improves to 0.40 and 0.29 when predicting the train and test data respectively when using shell and packing tissue volume ratios. A slight improvement is observed when allowing the model to use three explanatory variables, with mean *R*^2^ scores of 0.42 and 0.30 for predicting training and testing data respectively. The best trio consists again of shell and packing tissue volume ratio, and relative kernel density. Finally, we observe that adding more explanatory variables to the OLS does not improve the results. Moreover, the variability of *R*^2^ coefficients increases for the test data, which indicates that the model tends to overfit as more variables are taken into account (Figure 5A-B).

A similar analysis was performed on OLS linear models to predict shell strength, with shell volume ratio being the most predictive single trait. This produced an *R*^2^ coefficient of 0.53 and 0.49 if predicting with training and test data respectively. The variability of *R*^2^ coefficients is much lower than in the prior modeling of ease of kernel removal. The model can improve its *R*^2^ up to 0.66 and 0.62 respectively if it uses two explanatory variables instead, namely shell thickness and packing tissue volume ratio. The model shows no improvements, and actually performance worsens, when more explanatory variables are added to it, which indicates overfitting (Figure 5C-D). To predict kernel weight ratio, using simply the shell volume airless ratio, the model reports an average *R*^2^ score of 0.75 and 0.73 with training and testing data respectively. The variability of *R*^2^ is even lower than in the previous two models. This performances improves up to 0.84 and 0.82 if the model uses two explanatory variables instead. The best pair consists of shell thickness and kernel airless volume ratio. No improvements are reported whenever the model employs more explanatory variables (Figure 5E-F).

Finally, we perfomed a PCA using only the 6 most predictive morphological phenotypes discussed above: shell thickness, shell and packing tissue volume ratio, and shell and kernel airless volume ratio. The first two PCs explain more than 85% of the variance. We observe that accessions follow a gradient based on their scores of traits of breeding interest. If all the 49 morphological phenotypes are considered instead, the first two PCs now only explain 50% of the total variance. As with the OLS models, this suggests that more phenotypes tend to muddle the overall picture. Similarly, the gradient of breeding values is less clear. We highlight that the Earliest Himalayan accession (85-023-2) has a very small kernel which is notoriously hard to extract. Nonetheless, it is morphologically average, even when limited to the most predictive phenotypes, as it lies close to the center of the PCA (Figure 6). This averageness is also observed when examining the z-scores of all morphological phenotypes for every accession (Figure S5).

**FIGURE 6.**
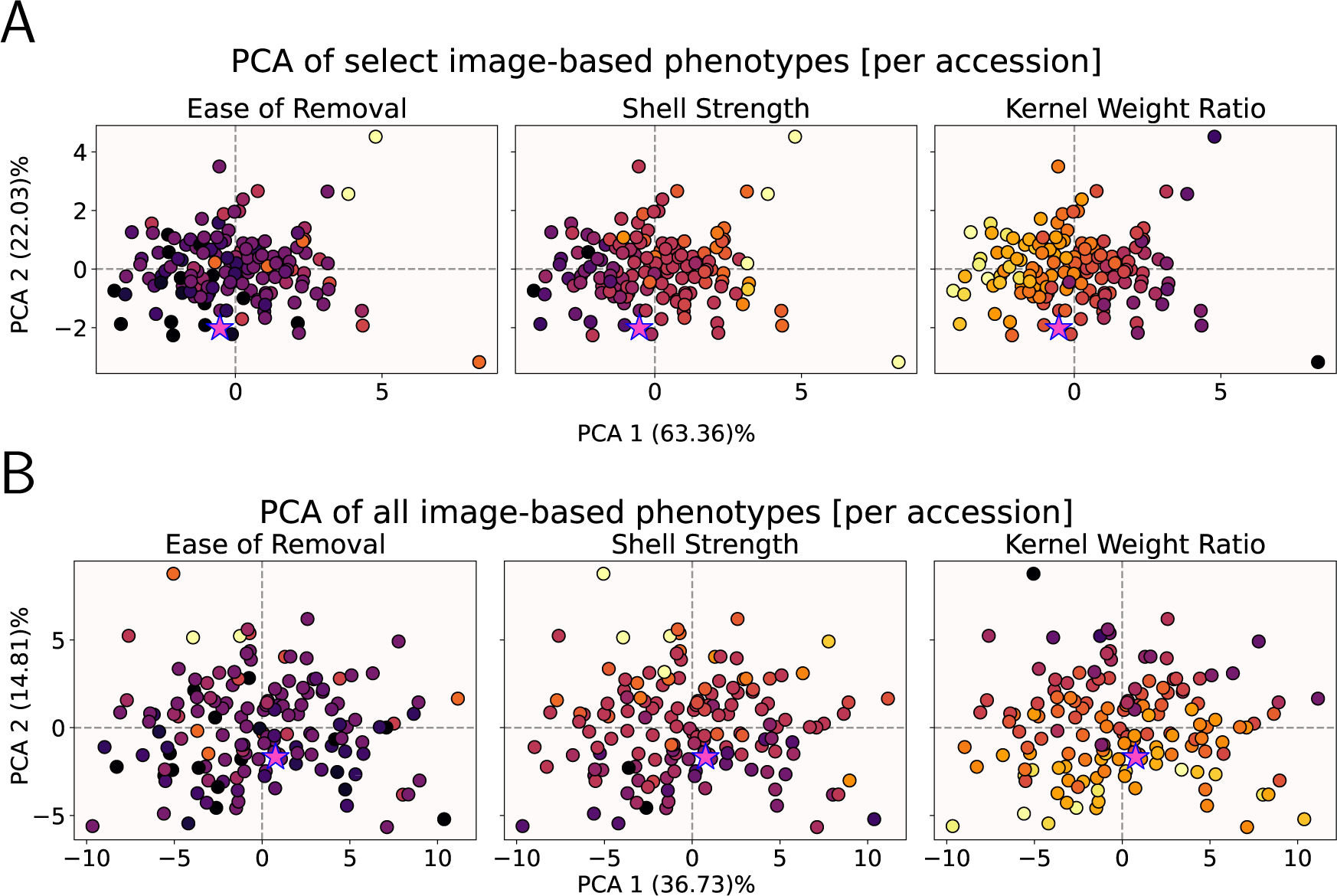
Principal Component Analysis using solely morphological phenotypes. The scatter plot is the same for each row, however the color varies depending on the trait of interest indicated on top. Darker colors represent lower values. The Earliest accession (85-023-2) is represented by a magenta star instead. (A) The PCA uses only the 6 most predictive morphological phenotypes as suggested by Figure 5. (B) The PCA uses all 49 morphological phenotypes from Table 3.

## 4 Discussion

There is plenty of observed phenotypic diversity within a fixed walnut population, but surprisingly these variations are usually not sufficient to distinguish geographically distant populations (Roor et al., 2017). This difficulty to comprehensively measure walnut morphology is more pressing when trying to understand the fine-grained details that determine important traits of commercial concern (Bernard et al., 2021). Three such traits are the ratio of kernel-to-total walnut weight, the kernel ease of removal —how easy is to remove its main two halves intact—, and shell strength. Multiple-pronged strategies have been proposed and developed to unravel the underlying mechanisms that regulate shell thickness, shape, and strength. Through genome-wide association studies (GWAS), transcription factors and pathways that affect the shell and seal formation have been identified (Sideli et al., 2020; Wang et al., 2022). Walnut shell physics have been explored with numerical simulations, where walnuts are modeled as thin spheres and biophysical mechanical properties are tested under unidirectional loads based on finite-element analyses (Bao et al., 2022; Koyuncu et al., 2004). Novel work has focused on the shape of the polylobate sclerid individual cells that tesselate and conform the walnut shell while forming intricate puzzles that confer remarkable toughness and strength (Antreich et al., 2019; Zhang et al., 2014). These tough puzzles in turn are determined by biochemical processes during walnut shell development, as individual cells go from a soft to hard state (Antreich et al., 2021).

X-ray CT scans allows us to accurately extract more nuanced shape and size features from our sampled walnuts, providing new avenues to explore subtle morphological changes and its implications. For example, with more size-related features, we can compute better allometric relationships that point to biophysical constraints in walnut growth development. For example, the growth of empty space within a walnut outpaces the overall nut growth rate, while the kernel growth rate falters (Figure 3). This indicates that larger walnuts tend to contain a higher proportion of air and a lower proportion of kernel, which agrees with past observations of inner air increase during walnut development Xiao et al. (2020). At the same time, we observed that for small nuts, their total volume was almost identical to the volume of their convex hulls. This implies that smooth, groove-free nuts must be small. Moreover, this allometric relationship only holds for nut that are larger than a certain size (16mm in diameter), which indicates that the growth dynamics of the nut undergo a regime change as the nut develops (Pinney and Polito, 1983; Zhao et al., 2016). All of the allometric observations and correlations above suggest that walnut and kernel sizes and smoothness are not just dependent on genes and environment, but there are also unavoidable biophysical constraints at play that should be explored further and considered by breeding programs (Niklas and Hammond, 2019).

Our extended list of measured phenotypes also offers new insight to qualitatively assessed traits. The biophysical constraints around air content appear to also play a role for several traits of interest. Walnuts with relatively low air content tend to be smaller in volume, have thicker and stronger shells, more packing tissue, the kernels are denser and harder to extract, with a lower weight ratio. The effects of air content can also affect correlation coefficients for other tissues. For example, if we consider air as part of the total walnut volume, then the relative volume of packing tissue is positively correlated to shell strength and ease of kernel removal. This agrees with reports that walnuts with thick packing tissue tend to be difficult to open (Kouhi et al., 2020; Mirmahdi and Khadivi, 2021; Sarikhani Khorami et al., 2014). These correlations vanish if we disregard air as part of the total walnut volume. However, new correlations emerge, as the airless ratio of packing tissue volume is now negatively correlated to shell thickness, volume, and density. Unlike packing tissue, the shell volume appears to be more independent from inner air content. Regardless if we consider air as part of the total walnut volume or not, we observe that walnuts with large shell volume ratio tend to have thicker, stronger, and denser shells, a low ratio of kernel volume and weight, and make kernel removal more difficult. This agrees with reports that thin shells allow easier kernel extraction (Amiri et al., 2010; Arzani et al., 2008; Fallah et al., 2022). Moreover, inner air may play a key role in walnut development. Examining the relative air volume compared to airless relative content of shell and packing tissue, the correlations suggest that as a walnut grows, packing tissue develops at the expense of shell tissue. This could be explained by the fact that the shell undergoes a careful biochemical balance between insulation and permeability of air and water as the walnut develops (Antreich et al., 2022) (Figure 4).

This nuanced relationship between air, shell, packing tissue during shell development is further highlighted when we consider their individual influence over ease of kernel removal. The single trait that can best explain and predict ease of removal is relative air volume. However, when the model is allowed to use any two morphological traits, the best results are produced when considering relative shell and packing tissue volumes, as opposed to relative air volume and something else. From a statistical point of view, the pair of shell-packing tissue volume ratios makes sense: both traits are completely uncorrelated and both are strongly correlated to air relative volume (Figure 4). This suggests that the influence of inner air into the ease of removal score can be understood as a combination of the relative shell and packing tissue volume influence (Figure 5A). We can make a similar argument for shell thickness and relative packing tissue volume encoding the same information as relative shell volume when it comes to understand each of these traits influence over shell strength (Figure 5C). We can also argue that the relationship between the kernel weight ratio and the shell airless volume ratio is already encoded by the kernel airless volume ratio and shell thickness. The inner air content, and the air ratio in a walnut are traits that often overlooked, yet they seem to play crucial roles in walnut development and appear to be intrinsically linked to traits of high commercial interest. Moreover, we can successfully encode commercially relevant scores using only volume- and thickness-based morphological traits (Figure 6). This opens a potential avenue for new phenotyping platforms focusing on just volumetric measurements.

Nonetheless, the exact link between broad morphological phenotypes and commercially relevant traits remains elusive. We highlight the Earliest accession (85-023-2) is morphologically average when compared to the general population as its corresponding z-scores are small for most of phenotypes (Figure S5D). Yet, its shell is notoriously strong and it is extremely difficult to extract its kernel intact. This accession is originary from the Himalaya region, which is one of the hotspots for walnut diversity (Aradhya et al., 2017) and the only wild accession that formed part of our study. This in turn poses exciting questions on morphological changes during walnut domestication.

Walnuts offer a especially unique opportunity to analyze domestication in perennial crops. Despite their long history with humans, current research suggests that walnut domestication happened less than 100 years ago (Mapelli et al., 2018). Even today, due to economical and horticultural reasons, walnut is mainly propagated via seeds instead of grafting throughout most of Southwest Asia (Rezaee et al., 2008; Thapa et al., 2021). This makes walnut an exciting organism to study the immediate effects of domestication and breeding in real time across multiple populations. We can especially focus on inner air, kernel, shell, and packing tissue volumes, as these can successfully encode other commercial traits. While X-ray CT scanning offers a powerful and precise way to quantify these volumes, there are inexpensive alternatives if we just want to measure volume. Kernel, shell, and packing tissue volumes can be measured via toluene (C_7_H_8_) displacement (Aydin, 2003; Gharibzahedi et al., 2012). Inner air volume can be estimated following the Archimedean’ principle and a vacuum flask partially filled with a surfactant solution (Marquard, 1989; Raskin, 1983). This is an exciting opportunity to design affordable phenotyping platforms. A careful, nuanced study of walnut morphology combined with extended low-cost phenotyping might provide us key insights into other domestication-induced phenotypical and genotypical changes, and accelerate the selection of progenitors in breeding programs.

## Acknowledgements

Daniel Chitwood is supported by the USDA National Institute of Food and Agriculture, and by Michigan State University AgBioResearch. The work of Elizabeth Munch is supported in part by the National Science Foundation through grants CCF-1907591, CCF-2106578, and CCF-2142713.

## Author contributions

EA, PB, EM, and DC conceived the experiment. PB selected the accessions to be scanned and collected the plant material, ensuring that walnut accessions were broadly represented. MQ and DC collected the digital data. EA developed the necessary scripts to process the scans and extracted their shape descriptors. EA analyzed the data and wrote the manuscript. All authors contributed, reviewed, and revised the manuscript.

## Software and data availability

The processed and cleaned walnut X-ray CT 3D reconstructions can be found in the Dryad repository https://doi.org/10.5061/dryad.ngf1vhj09, along with their separated tissues.

All our code in the form of python jupyter notebooks is available at the https://github. com/amezqui3/walnut_tda repository. This includes the image processing pipeline to clean the raw scans and segment the walnut tissues, the computation of all the evaluated phenotypes.

## Conflict of interests

None declared

## Supplementary data

**FIGURE S1.**
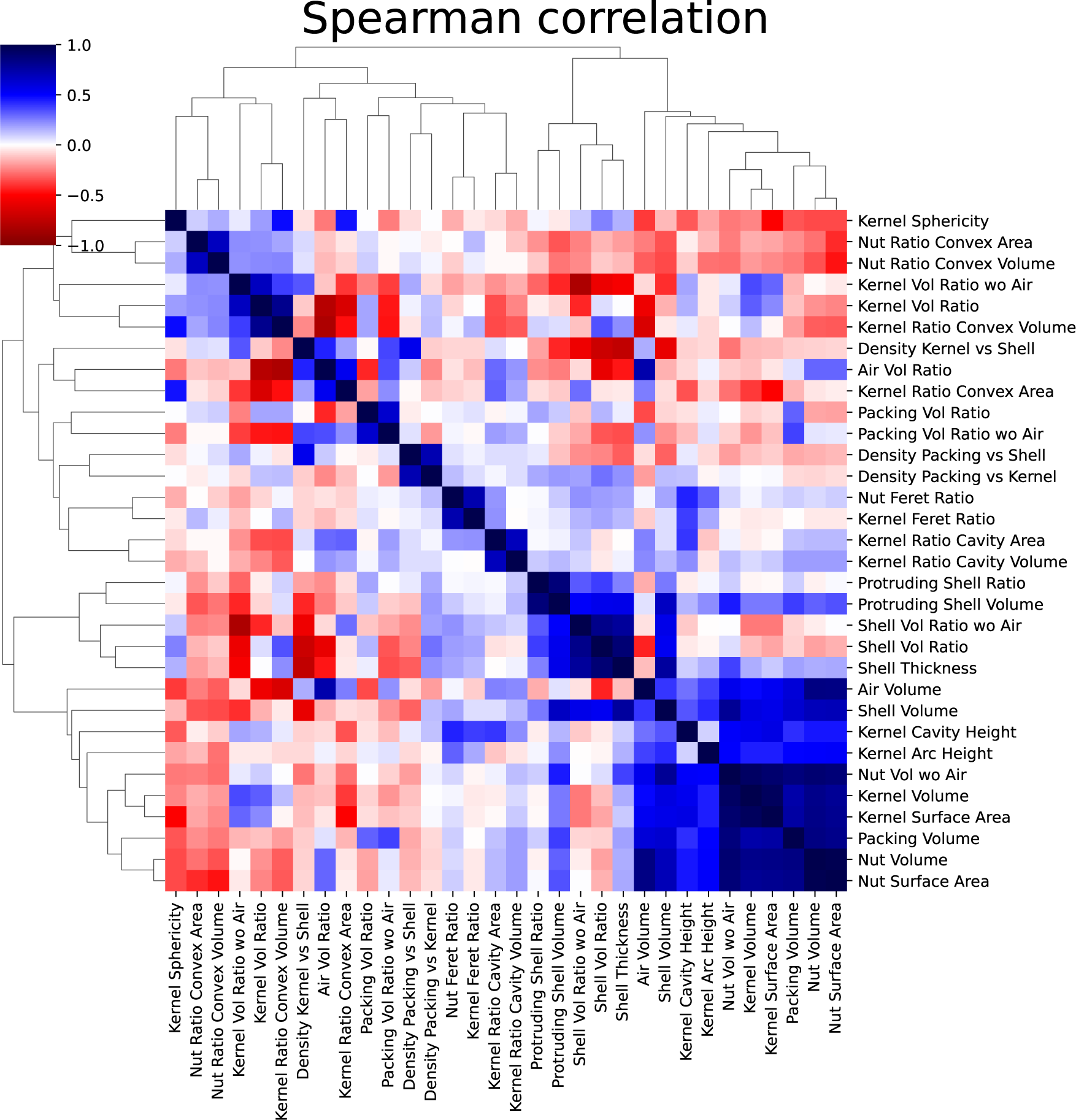
Spearman correlation for all phenotypes. Most of the overall nut size-related traits are only positively correlated with kernel size-related traits as expected. All the shell-specific traits are only positively correlated between themselves. All the sphericity, aspect ratio, and rugosity are highly correlated only among themselves.

**FIGURE S2.**
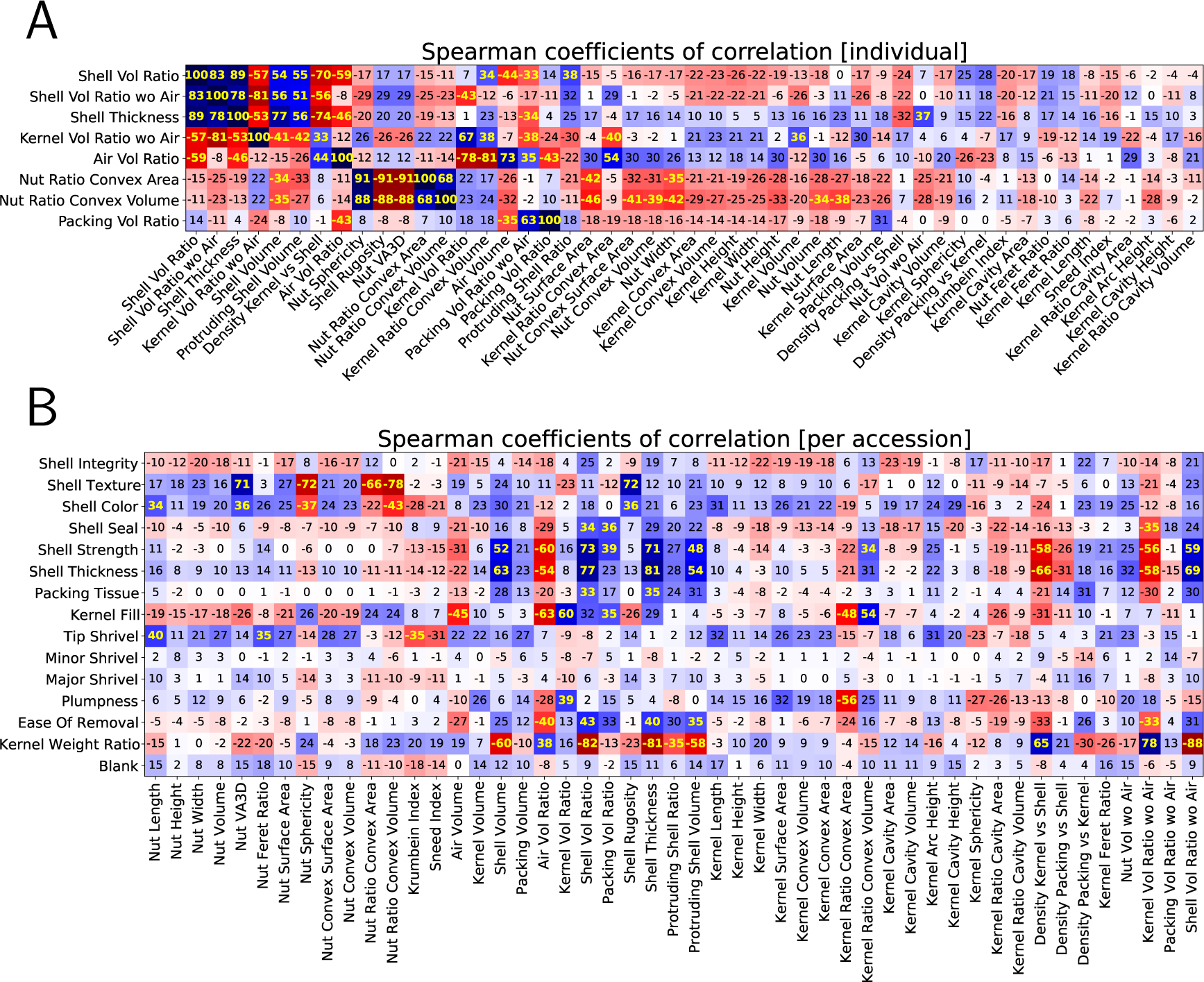
Detailed Spearman correlation indices for more pairs of traits. Indices have been multiplied by 100. Indices in yellow are highly significant (p-value *>* 0.001) with an absolute value larger than 0.35. (A) Correlations between select morphological phenotypes with all the 49 traits. (B) Correlations between all the traits of breeding interest and the 49 morphological phenotypes.

**FIGURE S3.**
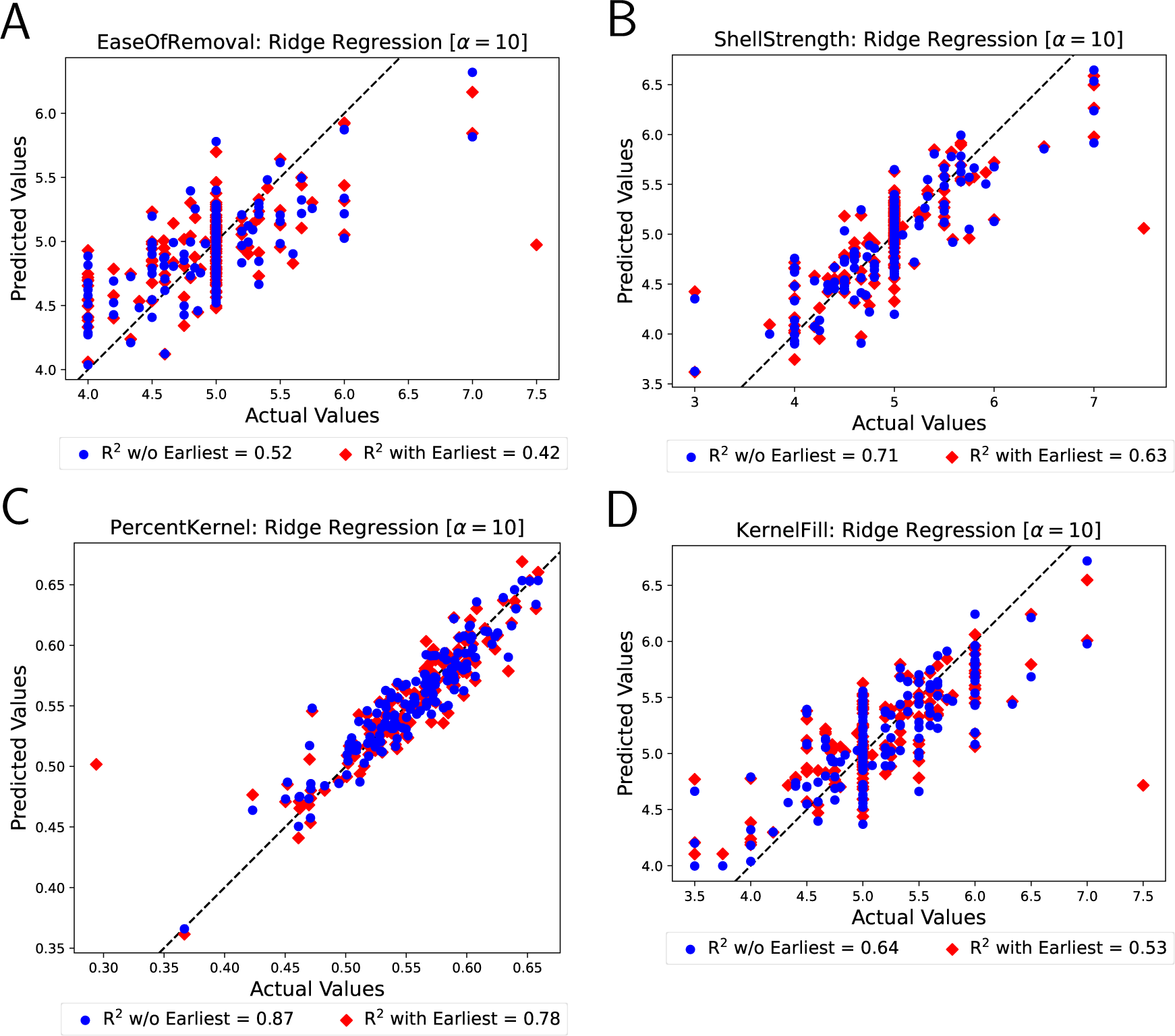
Ridge regression using all the 49 morphological phenotypes as descriptive variables to predict (A) Ease of Removal, (B) Shell Strength, (C) Kernel-to-total weight ratio, and (D) Kernel fill score. For all cases, we used an *α* = 10 regularization value for the ridge regression. Individual points represent individual accessions. Red diamonds represent predicted values when considering all 149 accessions, while blue circles are predicted values when excluding the Himalayan Earliest accession (UCACCSD 85-023-2). The *R*^2^ coefficient of determination is indicated at the bottom of each plot. Notice that excluding the Earliest accession improves considerably the *R*^2^ score.

**FIGURE S4.**
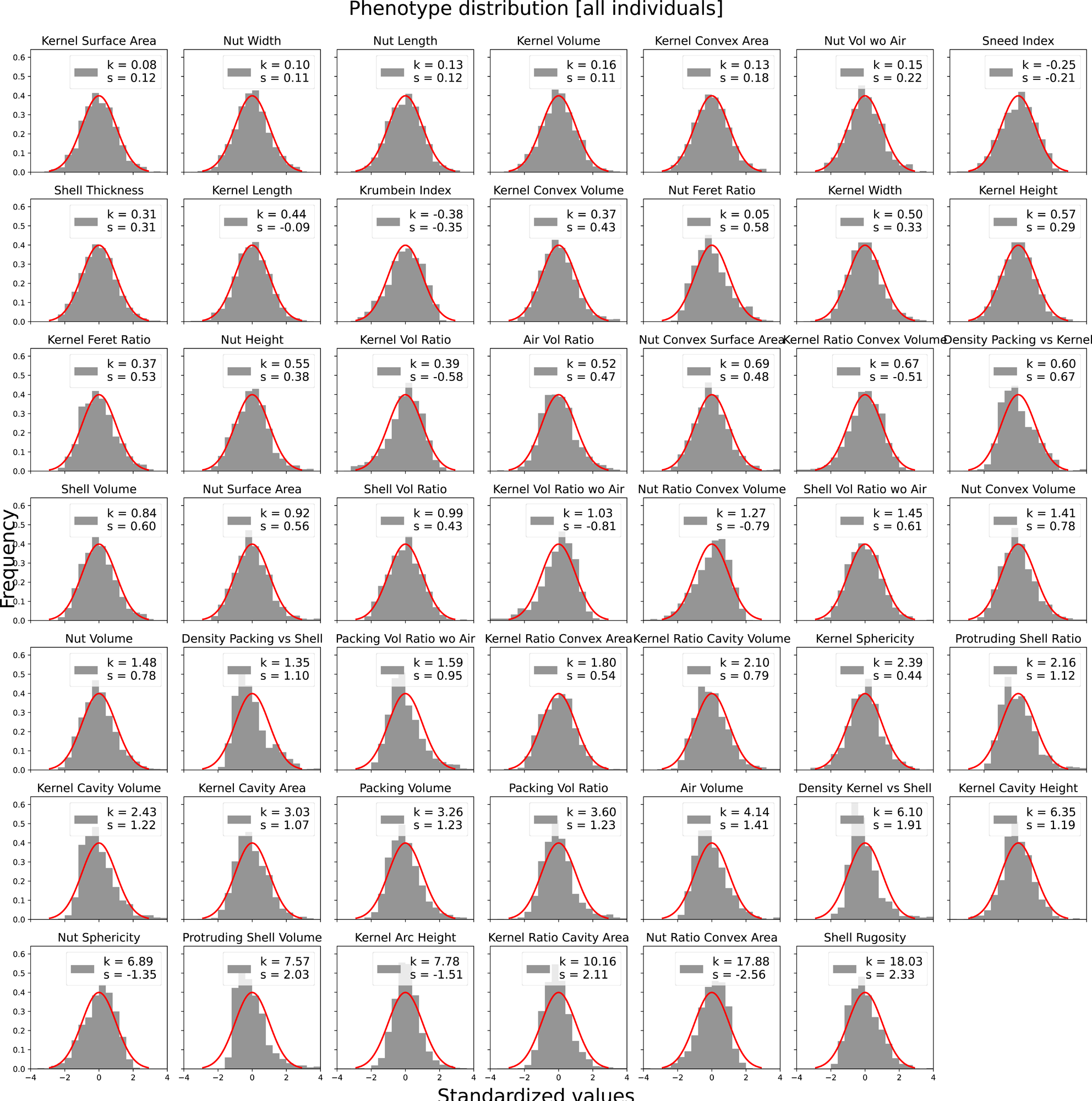
The values were centered at 0 and rescaled to unit variance. A normal Gaussian bell is drawn in red for comparison. The Fisher-corrected kurtosis (k) and skewness (s) are indicated for each trait distribution. For reference, a normal distribution has both *k* = 0 and *s* = 0.

**FIGURE S5.**
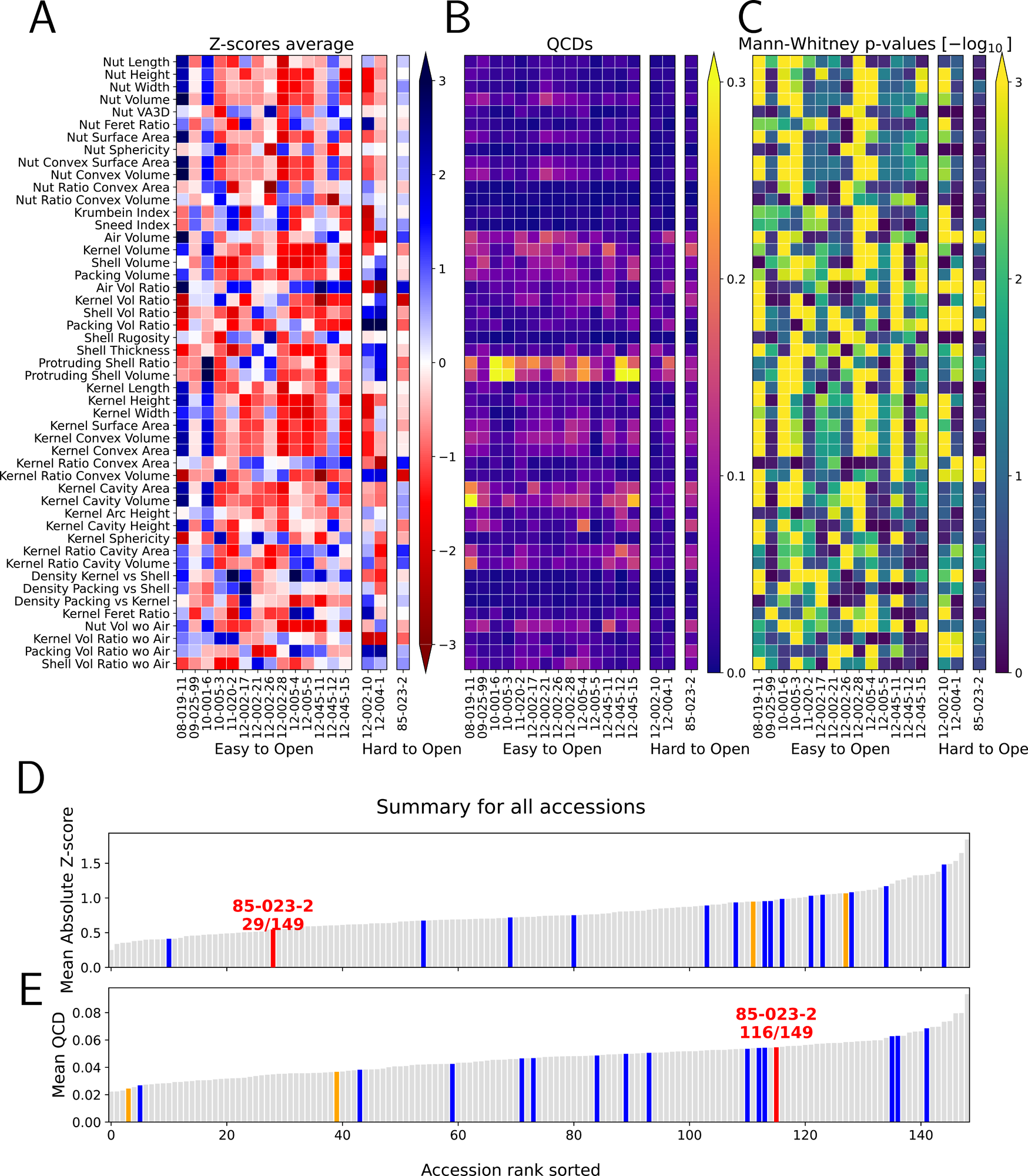
Variability and uniqueness of morphological traits for every accession. For every measured phenotype, we computed the mean and QCD values for individuals of the same accession. (A) Since most of the morphological phenotypes follow a normal distribution (Figure S4), the z-score of these means was computed. A large z-score would suggest that the accession shows a very different phenotypical value compared to the rest of the population. We were particularly interested to explore the trait distribution for accessions associated with the lowest and highest ease of kernel removal scores. (B) The QCD of the traits for individuals in the same accession quantifies how variable is every shape and size parameter within individuals of the same accession. (C) Mann-Whitney U-tests were performed to determine if certain morphological traits were sufficiently different to tell apart one accession from the rest of the population. (D) The mean absolute z-score provides an overall estimate on how distinctive are the morphological features of an accession. Accessions with the lowest ease of kernel removal scores are depicted in blue, while highest scores are in red and orange. (E) Similar to (C), the average QCD provides an overall estimate on how variable is the shape of individuals within the same accession.

## Notes

https://doi.org/10.5061/dryad.ngf1vhj09

https://github.com/amezqui3/walnut_tda

### Competing Interest Statement

The authors have declared no competing interest.

