## Supplementary figures and images for "The shape and volume of air, kernels and cracks, in a nutshell"

### Figure S1

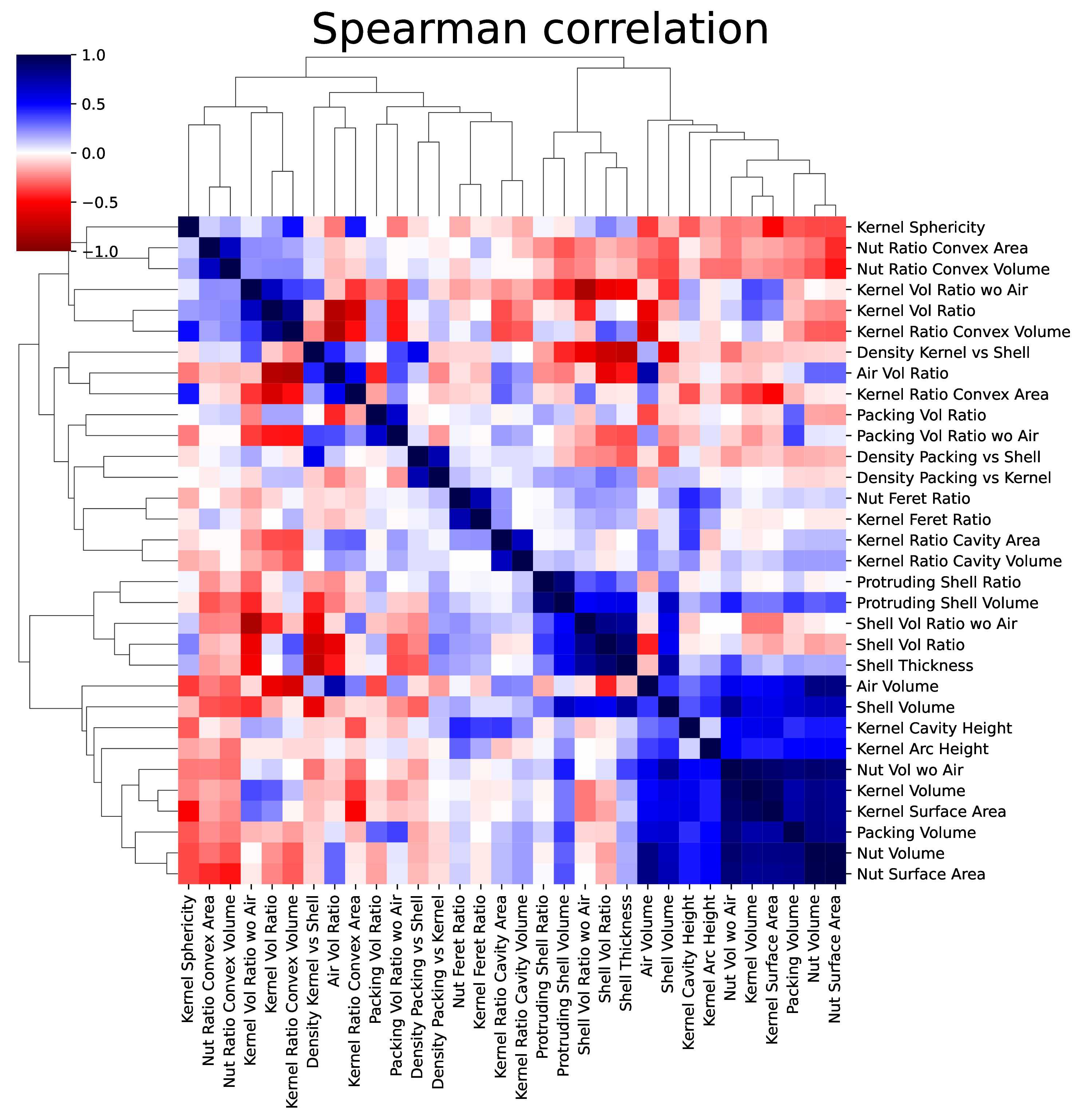

### Figure S2

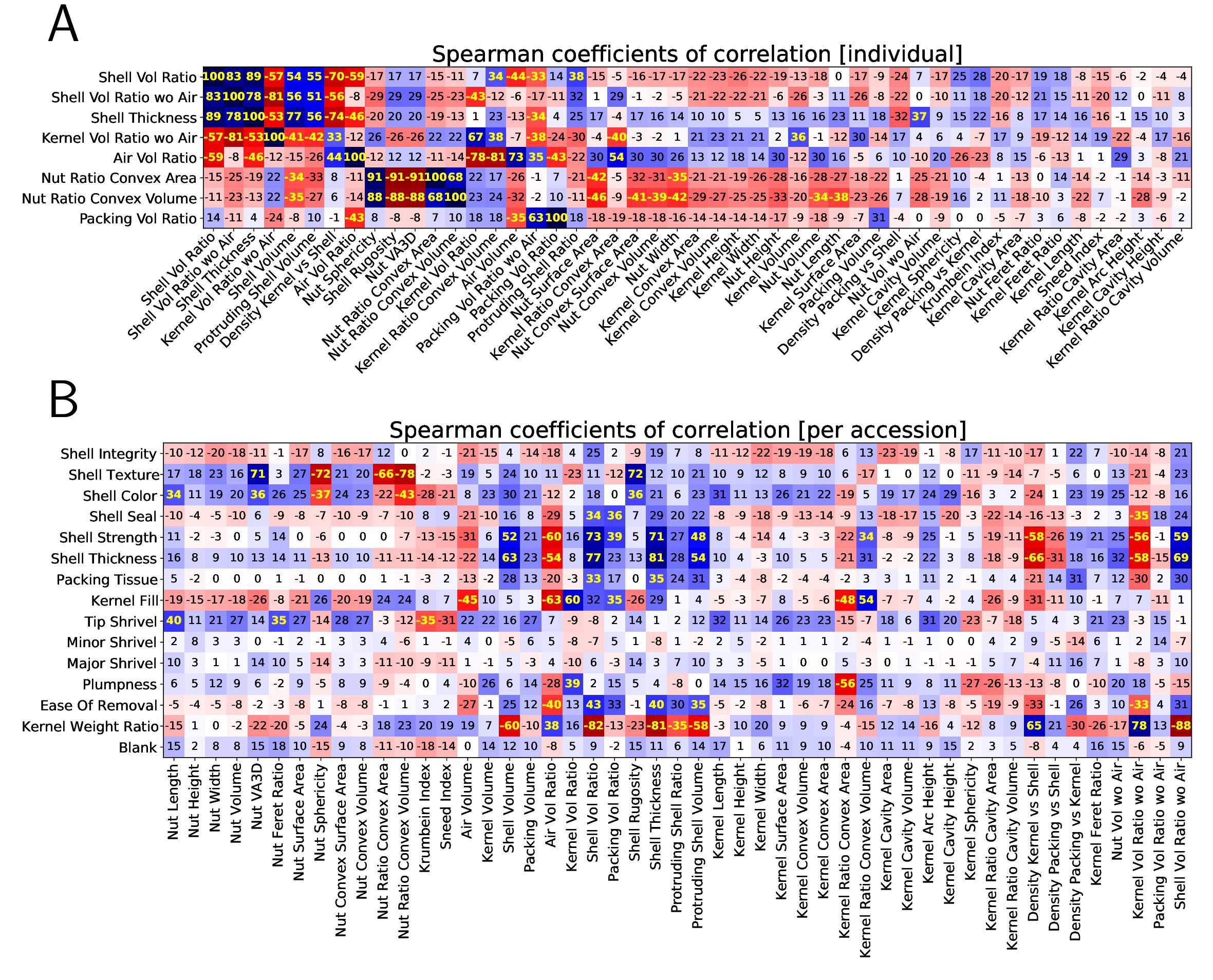

### Figure S3

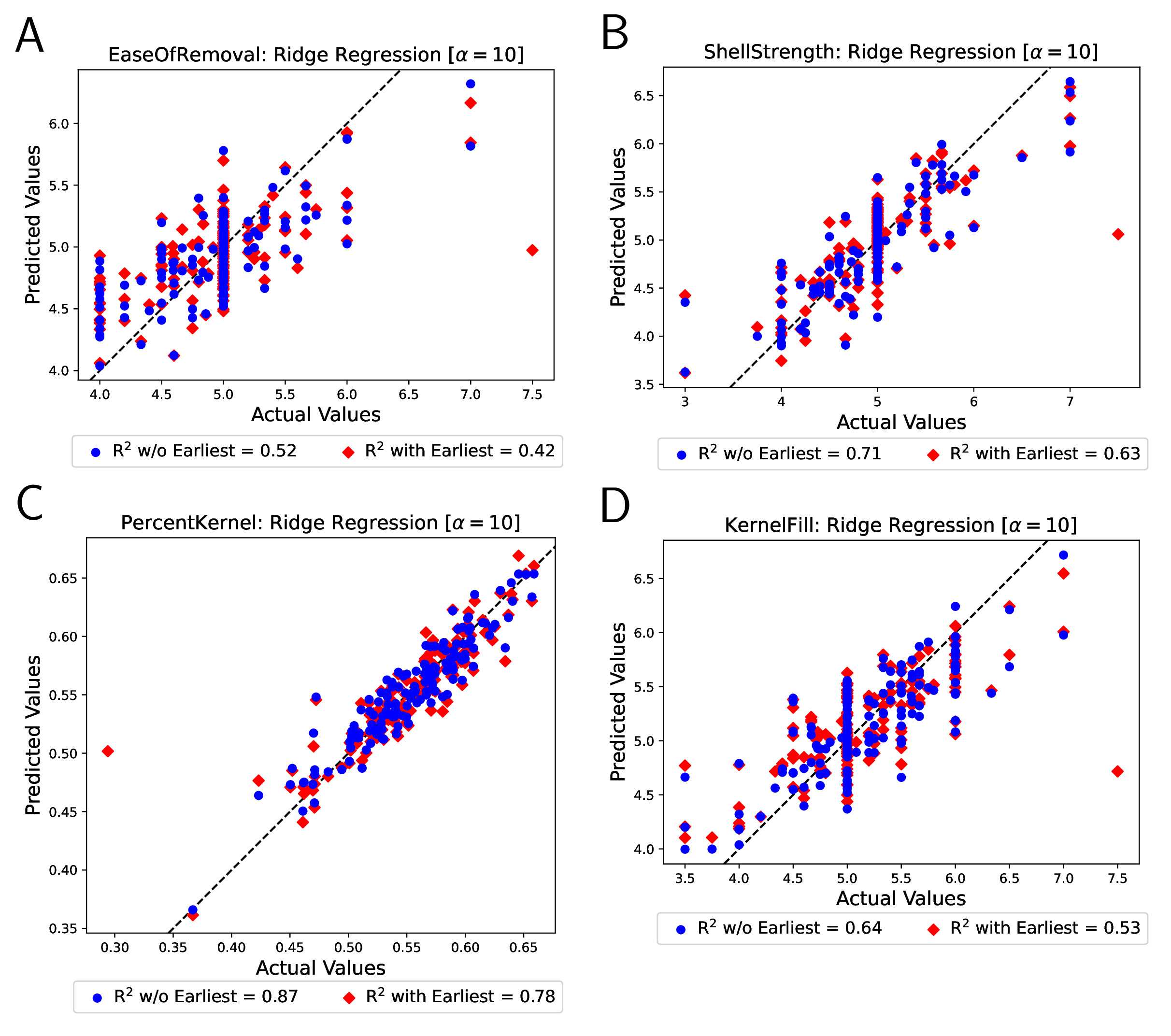

### Figure S4

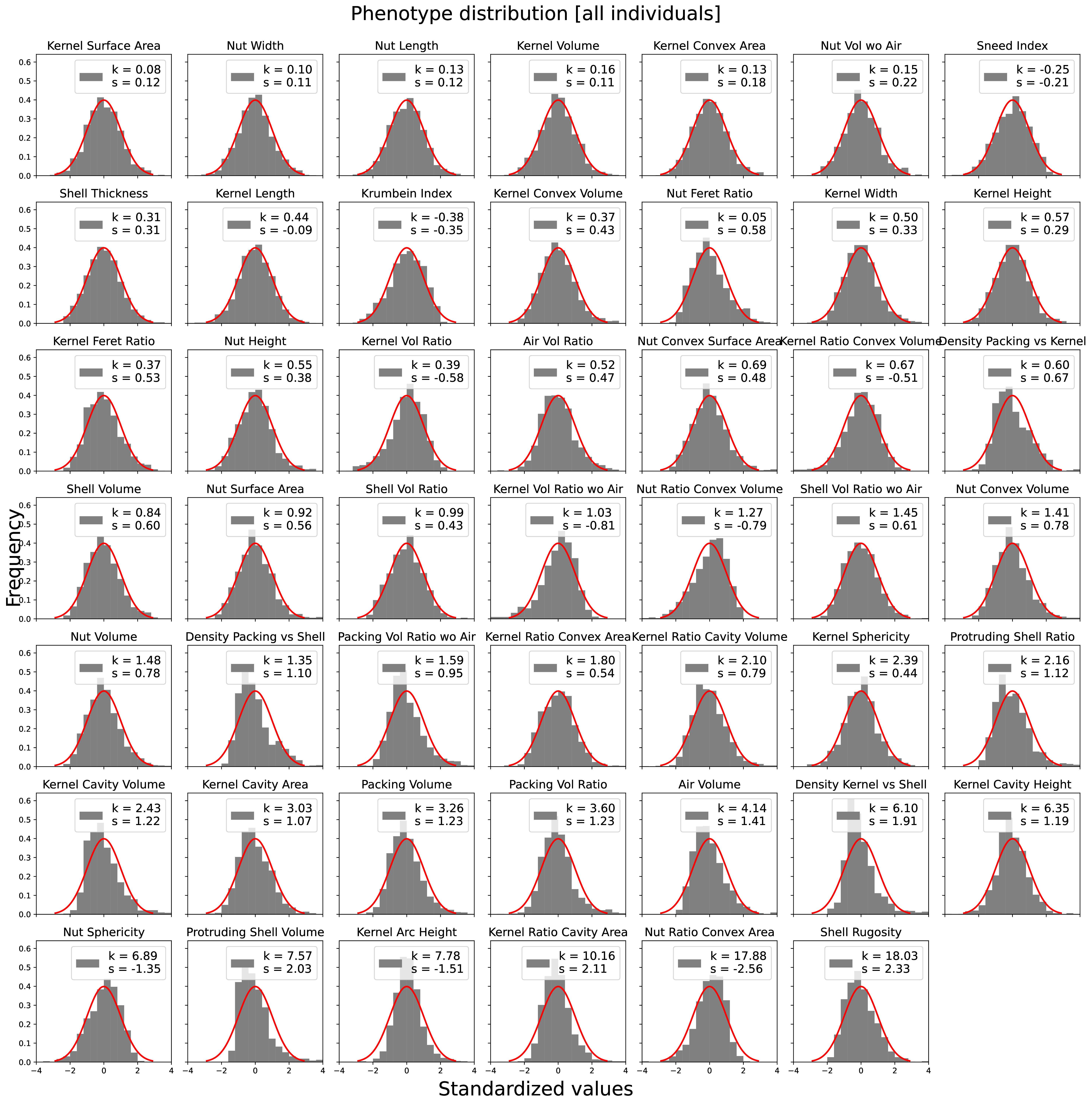

### Figure S5

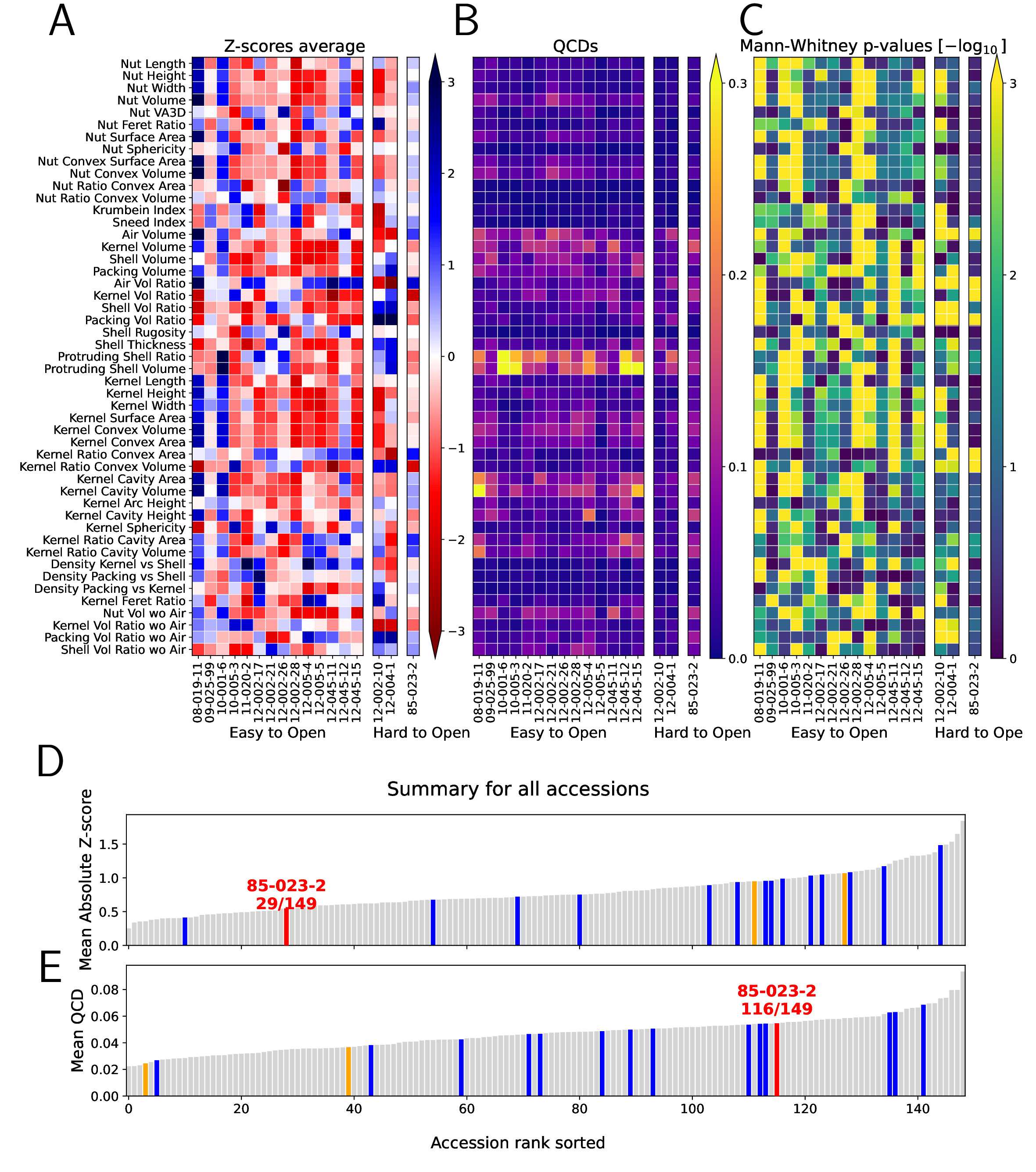
